# Circulating levels of epirubicin cause endothelial senescence while compromising metabolic activity and vascular function

**DOI:** 10.1101/2020.06.25.171249

**Authors:** Amanda J Eakin, Tamara McErlain, Aileen Burke, Amy Eaton, Nuala Tipping, Gloria Allocca, Cristina M. Branco

## Abstract

Anthracycline-based chemotherapy is a common treatment for cancer patients. Because it is delivered intravenously, endothelial cells are exposed first and to the highest concentrations, prior to diffusion to target cells. Not surprisingly, vascular dysfunction is a consequence of anthracycline therapy. While chemotherapy-induced endothelial damage at administration sites has been investigated, the effects of lower doses encountered by distant microvascular networks has not. The aim of this study was to investigate the impact of epirubicin, a widely used anthracycline, on healthy endothelial cells to elucidate its effects on microvascular physiology.

Here, endothelial cells were briefly exposed to low doses of epirubicin to recapitulate levels in circulation following dilution in the blood and compound half-life in circulation. Both immediate and prolonged responses to treatment were assessed to determine changes in endothelial function.

Epirubicin caused a decrease in proliferation and viability in hUVEC, with lower doses resulting in a senescent phenotype in a large proportion of cells, accompanied by a significant increase in pro-inflammatory cytokines and a significant decrease in metabolic activity. Epirubicin exposure also impaired endothelial function with delayed wound closure, reduced angiogenic potential and increased monolayer permeability downstream of VE-cadherin internalization. Primary lung endothelial cells obtained from epirubicin-treated mice similarly demonstrated reduced viability and functional impairment. *In vivo*, epirubicin treatment resulted in persistent reduction in lung vascular density and significantly increased infiltration of myeloid cells.

Modulation of endothelial status and inflammatory tissue microenvironment observed in response to low doses of epirubicin may predict risk for long-term secondary pathologies associated with chemotherapy.

## Introduction

Chemotherapy is the standard of care for most cancer patients, especially those affected by aggressive cancer types, metastatic or refractory disease (NICE 2020). All chemotherapy is systemic, and mostly administered intravenously, at maximal tolerated doses (Le Tourneau, Lee et al. 2009). Even though cytotoxic drugs affect primarily their target cells (cancer cells), their effects are experienced and responded to also by somatic cells.

Anthracyclines are a family of commonly used chemotherapeutic agents, with demonstrated success in elimination of cancer cells and association with increased survival (EBCTCG 1988, EBCTCG 2005). Mechanistically, these drugs work by inhibition of topoisomerase II activity, compromising DNA and RNA synthesis, by intercalating nucleic acid chains (Szuławska and Czyz 2006), which results in cell death, primarily in rapidly dividing cells, with higher DNA replication frequency.

Doxorubicin is the most widely used anthracycline, used in the treatment of malignancies such as breast cancer, bladder cancer, or lymphoma (EBCTCG 2005, Gianni, Norton et al. 2009, Lori, Stein et al. 2010, Fukuokaya, Kimura et al. 2020). The cancer treatment effectiveness of Doxorubicin is however, accompanied by severe cardiotoxicity, and as a result it has been increasingly replaced by epirubicin, an epimer associated with milder adverse effects (Khasraw, Bell et al. 2012).

With increased survival, the long term consequences of anthracycline treatments have become more apparent, and beyond myocardial dysfunction, post-treatment conditions are now known to include hypertension and thrombosis, ultimately resulting in a poorer quality of life (Swain, Whaley et al. 2003, Jones, Haykowsky et al. 2007, Mercuro, Cadeddu et al. 2007, Barrett-Lee, Dixon et al. 2009, Cameron, Touyz et al. 2016). The usual administration by intravenous route, from which it reaches target cells by diffusion, inevitably exposes endothelial cells (EC) to the highest drug levels, even in organs distant from infusion site, and not surprisingly a severe consequence of anthracycline treatment is vascular dysfunction (Cameron, Touyz et al. 2016).

Vascular EC are critical in maintaining the tissue microenvironment as they create a barrier, a single cell layer that is both continuous and heterogeneous, between circulating blood and surrounding tissue. They are also responsible for local control of perfusion to match tissue demand and supply (Augustin and Koh 2017), and are the first responders to systemic signals of both internal or external nature, plastically adjusting flow and permeability to ensure organ homeostasis.

The damage inflicted by chemotherapeutical agents at injection sites has been investigated (Sato, Kondo et al. 2014). Specifically, a recent study found an association of Doxorubicin treatment with increased EC death, in which a significant inflammatory response, downstream of increased levels of pro-inflammatory cytokines and mediated by activation of the NFĸB pathway was also shown (Sonowal, Pal et al. 2018). Doxorubicin has also been associated with senescence of somatic cells, including EC, resulting in reduced regenerative ability, likely behind disruption of endothelial plasticity (Yasuda, Park et al. 2010, Wojcik, Buczek et al. 2015, Cappetta, Rossi et al. 2018).

The mechanisms underlying these observations and the impact on co-morbidity associated with cancer treatment are not well understood, but effects on microvasculature in particular can underlie severe and persistent effects on organ function. Importantly, most studies to date explore damage of injection sites (high drug concentrations) (Yamada, Egashira et al. 2012), and not the low level, sublethal doses encountered by the majority of somatic cells. Responses and adaptation of EC to cytotoxic exposure may compromise the organs in which they reside, including dysfunction, degeneration or indeed potential for onset and success of distant metastases (Kennecke, Yerushalmi et al. 2010, Martin, Cagney et al. 2017, Xiao, Zheng et al. 2018).

The aim of this study is to investigate the consequences of low-level exposure of epirubicin on healthy EC. In spite of its known reduced toxicity in comparison to Doxorubicin, this drug has not been studied in the context of its effects on microvascular physiology and the downstream impact this may have on vascular pathogenesis.

In the present study, EC were exposed to short treatments of physiological doses of epirubicin, to emulate exposure in following drug dilution in the blood stream, and accounting for its relatively short half-life (Robert 1994). The aim was to investigate how EC response may educate the organ microenvironment, to what extent, and the duration and/or reversibility of EC adaptations.

Human umbilical vein EC (hUVEC) were used to investigate effects on EC survival and function. Additionally, naïve (tumour-free) mice were treated with intravenous epirubicin to assess effects of treatment on lung vascular parameters, considering that lung disease and associated comorbidities are heavily associated with vascular dysfunction (Reiterer and Branco 2020), and this is also a common metastatic site. Primary microvascular EC (MVEC) were also isolated from lung tissue (Reiterer, Colaco et al. 2019) to investigate the effects of epirubicin on EC behavior following *in vivo* exposure.

## Materials and Methods

### Cell culture

hUVEC were maintained in collagen-coated plates, at 37 °C and at physiological oxygen tensions (5 % O_2_) and 5 % CO_2_. Cells were expanded in antibiotic-free medium containing a 1:1 mix of Ham’s F12 nutrient mix (21765037, Gibco) and low glucose Dulbecco’s Modified Eagle’s Medium (DMEM) (D6046, Sigma Aldrich), supplemented with 1 % MEM non-essential amino acids (11140050, Gibco), 2 mM sodium pyruvate (11360070, Gibco), 20 mM Hepes (H0887, Sigma-Aldrich), 10 mg/mL Heparin (H3149-25kU, Sigma-Aldrich), 7.5 mg/mL EC growth supplement (E2759, Sigma-Aldrich) and 20 % fetal bovine serum (F7524, Gibco).

### Migration assay

Cells were grown to > 90 % confluence in a 12-well plate and incubated with saline or epirubicin at concentrations typically found in circulation 3 μg/mL or 5 μg/mL for 30 min followed by replacement with drug-free medium. 10 μg/mL is typical but proved lethal for hUVEC *ex vivo*. Cells were analyzed immediately (baseline) or allowed to recover for a given period before beginning the assay. Prior to the migration assay, growth medium was replaced by serum-free basal medium containing 50 μg/mL Mitomycin C (CAY11435, Cambridge Bioscience) for 1.5 h. This was removed at the start of the assay and a scratch/wound was made by scraping a 1 mL pipette tip through the middle of each well. Cells were washed twice in PBS before adding growth medium, and images taken at various time points throughout the assay. Wound closure was quantified using ImageJ software (version 8.0).

### Angiogenesis assay

10 μL of complete Matrigel (354234, Corning) was loaded onto angiogenesis slides (1B-81506, Thistle scientific) and incubated for at least 30 min at 37 °C to solidify the matrix. > 90 % confluent cells grown in 12-well plates were treated, as before, with epirubicin or saline for 30 min. 4.5 × 10^3^ cells in 50 μL of growth medium were seeded on top of the Matrigel substrate. Images were taken at various time points throughout the assay. Tube formations were analyzed using the ‘Angiogenesis Analyzer’ plug-in for ImageJ.

### Permeability assay

hUVEC were plated at a density of 4×10^4^ cells/well into 8 μm Fluoroblok™ inserts (351152, Appleton Woods LTD) in a 24 well companion plate (353504, Analab). Cells were also plated at the same density into a 96-well plate (which have the same surface area as the Fluoroblok™ inserts) to monitor confluency; when the cells formed a uniform monolayer in the 96-well plates, they were assumed confluent in the inserts, and at that time incubated with saline or epirubicin for 30 min followed by replacement with drug-free medium. Cells were either assayed immediately after treatment or allowed to recover for 1 d. 1 mg/mL FITC-Dextran 70 kDa (46945, Sigma-Aldrich) in growth medium was added to the upper chamber, and medium without FITC-Dextran was added to the lower chamber immediately before the start of the assay. Fluorescent signal (488 nm) was read every 5 min for a total of 6 h using an Omega plate reader (BMG Labtech) set at 37 °C, 5 % O_2_ and 5 % CO_2_. Signal was normalized to positive control wells (FITC dextran was added to the lower chamber for maximum fluorescence signal), and negative controls (no FITC-Dextran was added to either chamber); assay also included a no-cell control.

### Metabolism assays

Local extracellular acidification rate (ECAR) and oxygen consumption rate (OCR) following cytotoxic therapy in hUVEC were measured using a Seahorse XFe96 Analyzer (Agilent) and compared to saline controls. 1×10^4^ hUVEC grown at 5 % O_2_, 5 % CO_2_ at 37 °C were seeded on Seahorse microplates pre-coated with 3 μg/mL collagen I (C9791, Sigma Aldrich) in 0.1 M acetic acid and allowed to adhere for at least 12 h. hUVEC were treated with 3 μg/mL epirubicin for 30 min. Medium was replaced with drug-free medium and cells were analyzed immediately (baseline) or after recovering for 1 d. The day before the assay was carried out, Seahorse XF assay Medium was adjusted to a pH of 7.4 and supplemented with 2.5 mM glutamine for the glycolytic stress test, and with 2.5 M glucose and 100 mM pyruvate for the mitochondrial stress tests. The injection cartridge was hydrated overnight in a CO_2_ free incubator at 37 °C. To carry out the analysis at non-atmospheric conditions, the Analyzer was placed in a Ruskinn hypoxia chamber and atmosphere equilibrated to 5 % O_2_ and 0.1 % CO_2_. The instruments, Seahorse XF Medium (103575, Agilent) and calibrant solution were also equilibrated to the adequate oxygen levels for a minimum of 12 h prior to the assay. Solutions for injections were prepared fresh immediately before each assay. For the glycolytic stress tests, a final concentration of 10 mM glucose, 1 μM oligomycin and 50 mM 2-deoxyglucose were used. For the mitochondrial stress tests, compounds were added at final concentrations of 1 μM oligomycin, 0.5 μM FCCP and 0.5 μM Antimycin A/Rotenone. A final volume of 200 μL was kept consistent across test and control wells, with vehicle injections added to negative controls to allow volume normalization throughout the assay. Each measurement followed a 25 min equilibration period to establish the baseline levels, and was composed of a 5 min mix- 0 min – 2 min measure cycle which lasted as recommended by the manufacturer. All Seahorse measurements were normalized to the μg of total protein concentrations in each well, quantified with Bicinchoninic acid (BCA) assay (23225, ThermoFisher). Seahorse Wave software and GraphPad were used to statistically analyze these data.

### TUNEL assay

The TUNEL assay was performed using the Invitrogen by Thermo Fisher Scientific Click-iT™ TUNEL Alexa Fluor™ 488 Imaging Assay (C10245, Thermo), as per the manufacturer’s instructions. hUVEC were seeded onto collagen-coated glass chamber slides and maintained until they reached 80 % confluence, with medium changed daily. hUVEC were treated with epirubicin for 30 min as described above. Slides were washed in PBS then fixed with pre-cooled acetone for 7 min followed by 3 washes in PBS. The cells were permeabilized with 0.4 % Triton-X in PBS for 20 min at room temperature. A 10 min incubation with TdT reaction buffer at room temperature was followed by a 1 h incubation at 37 °C in a humidified chamber with the TdT reaction cocktail (TdT reaction buffer, EdUTP and TdT). The slides were then washed twice in PBS containing 3 % BSA, and the Click-iT reaction cocktail (Click-iT reaction buffer and additive) was added to the slide chambers and incubated for 30 min at room temperature, protected from light, and finally washed three times for 5 min in PBS containing 3 % BSA. Nuclei were stained by incubation with 2 μg/mL Hoechst 33342 for 15 min at room temperature, protected from light. After two 5 min washes in PBS, slides were mounted with Vectashield Antifade mounting medium (H-1000, Vectorlabs) and imaged the following day using a Leica DM5500 fluorescent microscope.

### Senescence assay

The senescence assay was performed using the Abcam beta Galactosidase staining kit (ab102534, Abcam) as per the manufacturer’s instructions. hUVEC were plated onto collagen coated 12-well plates and grown until 80 % confluence. Following epirubicin treatment, hUVEC were washed briefly in PBS before addition of the fixative solution (provided) and incubated for 15 min at room temperature. The cells were washed twice with PBS before addition of the staining solution (staining solution, staining supplement and X-Gal in DMSO). The plate was covered and incubated at 37 °C overnight inside a sealed bag to prevent any pH changes as a result of CO_2_ levels in the 37 °C incubator that may affect color development. The cells were washed briefly in PBS the following day and were counterstained with Nuclear Fast Red (N3020, Sigma-Aldrich) for 5 min then washed twice in PBS before imaging.

### Transcription factor assay

Transcription factor activation was assessed using the Signosis TF Activation Profiling Plate Array I (FA-1001-NF, Signosis). hUVEC were grown in 6-well plates until full confluence was reached. Wells were treated as before in triplicate with either saline or 3 μg/mL epirubicin. Following incubation, cells were left for 1 d in complete growth medium. The supernatants were removed, and nuclear protein was extracted using the Signosis Nuclear Extraction kit, according to the manufacturer’s instructions (SK-0001, Signosis). Nuclear protein was quantified using a BCA assay, as above, and 15 μg of nuclear protein per condition was used for the assay. The assay was carried out according to the manufacturer’s instructions, and results quantified with an OMEGA plate reader (BMG Labtech).

### Cytokine screening

Cytokine analysis was carried out using the Biotechne Human Cytokine Array (ARY005B, Biotechne). hUVEC were grown in a 6-well plate until confluent. Wells were treated as before, in triplicate, with either saline or 3 μg/mL epirubicin. Following treatment, cells were left to recover for 1 d in drug-free medium. The supernatants were removed and frozen at −80 °C until required for the assay. The assay was carried out by incubating neat media with pre-probed membranes (provided), washed and detected according to the manufacturer’s instructions. Blots were imaged using a Syngene image analysis system with GeneSys software, and signal was quantified and analyzed using ImageJ.

### Real-time polymerase chain reaction (qPCR)

RNA was extracted from fast-frozen epirubicin-treated hUVEC using a Purelink RNA mini kit (12183018A, ThermoFisher) followed immediately by a DNAse Treatment using an RNAse-free DNAse kit (79254, Qiagen). RNA was quantified using a Nanodrop and a minimum of 200 ng was used for reverse transcription cDNA synthesis. cDNA synthesis was performed using the Roche Transcriptor First Strand Kit (04896866001, Roche). qPCR was carried out using cDNA diluted 1:7, the Roche LightCycler 480 SYBR green I Master (04707516001, Roche) and Quantitect primers; β-Actin (QT00095431), VEGFA (QT01682072) and NOS3 (QT00095431). Target signal is normalized to β-Actin and presented as average fold change (treatment/control) ± SEM.

### Animal experiments

The study was conducted within the ethical principles of the Animal Welfare Act 1986, approved by the UK Home Office and the institutional ethical committee. C57BL/6 mice (aged 6-8 weeks) received 150 μL of 2 mg/mL epirubicin or saline control by intravenous route (tail vein). This can be considered a sub-clinical dose of epirubicin, approximating at 15 mg/kg (for a 20 g mouse), compared to an average clinical dose of approximately 30 mg/kg (U.S. Department of Health and Human Services 2005). Lungs were collected from one group of animals 1 d following injection to assess immediate effects, or after 7 d, to assess persistence or reversibility. A third group of animals received two injections, one on day 0 and one on day 3 to assess cumulative effects. Treatment groups had either 5 animals (1 d and 7 d re-injected) or 4 animals (7 d, one injection), and the same number of control animals per group, injected with sterile saline. Lungs were inflated with 10 % formalin and maintained in fixative overnight. Following multiple washes in PBS, tissue was embedded in paraffin for sectioning and immunofluorescent staining. Data from saline groups were ultimately combined and presented as one control group, following confirmation that there were no differences between the three saline-treated groups.

### Murine endothelial cell (EC) isolation and culture

Primary microvascular EC (MVEC) were isolated from lungs on day 1 and day 7 following treatment to assess proliferation, viability and function *ex vivo.* Lung tissue was removed following cervical dislocation, and stored in cold DMEM for up to 1 h until processing. Five mice were used per group. EC were isolated as previously described (Reiterer, Colaco et al. 2019) expanded and maintained at 37 °C, 5% CO_2_ and 10% O_2_ (physiological for lung EC) in the same medium as described for hUVEC (above). All functional assays were performed in triplicate, using cells at passage 1.

### Immunofluorescent (IF) staining

Paraffinized tissue from murine lung was sectioned (10 μm) using a microtome and mounted onto slides. Prior to staining, slides were deparaffinized in Histoclear II (H2779, Sigma-Aldrich) and rehydrated through graded alcohols. Acidic antigen retrieval was performed using a citrate-based buffer. After 3 washes in PBS slides were incubated for 10 min in 3 % H_2_O_2_ in methanol and blocked with Universal Protein Blocking Agent (GTX30963, Stratech) for 7 min. Slides were then incubated with Sudan Black (199664, Sigma Aldrich) for 10 min. After 3 washes in PBS, slides were incubated overnight at 4 °C in primary antibody (goat anti-mouse Podocalyxin (AF1556, R&D systems) for EC detection (Li, Li et al. 2001), rat anti-mouse Mac-2 (Galectin-3, 125402, Biolegend) for myeloid cells (Ho and Springer 1982), alpha smooth muscle actin – Cy3 (C6198, Sigma Aldrich) to stain pericytes (Babai, Musevi-Aghdam et al. 1990)). Slides were washed 3 times in PBS and then incubated for 2 h in secondary antibody (1:200 dilution AlexaFluor647 conjugated donkey anti-goat (ab150135, Abcam) and 1:200 dilution FITC-conjugated rabbit anti-rat (ab6730, Abcam)) protected from light. Finally, slides were incubated with 2.5 μg/mL 4’,6-Diamidino-2-Phenylindole, Dihydrochloride (DAPI) (D1306, Thermo) for 10 min and mounted with Vectashield Antifade mounting medium (H-1000, Vectorlabs). Imaging was performed the next day, using a Leica DM5500 fluorescent microscope at 20 × magnification. Eight non-consecutive sections were stained per animal, and 3 images were taken per section, subsequently analyzed using ImageJ software. Background correction was performed across all images by subtracting background with a rolling ball radius of 50 pixels. The plug-in ‘Vessel Analysis’ was used to quantify the vascular density and vessel diameter. The vascular density was quantified using the green color channel and enhancing local contrast to block size = 9; max slope = 4.00. The image was then made binary and the *vascular density* function. Average vessel diameter was determined by enhancing local contrast to block size = 9; max slope = 4.00, then adjusting the threshold to black and white and using the *diameter measurements* function. Pericyte to EC ratio was measured by adjusting the threshold to black and white and setting measurements to ‘Area Fraction’ and ‘Limit to Threshold’ before measuring. This was repeated for both color channels, then the pericyte signal was divided by the EC signal and the average taken per section. Macrophages were counted both by hand and using the particle analysis function on ImageJ. Both methods produced similar results and so the automated method was used for efficiency. The parameters set for particle analysis were 1) a size of greater than 80 pixels and 2) a circularity of between 0.45-1.00. Following analyses of all parameters, no differences were seen between the three saline control groups, those were pooled when plotted against the treatments (figures 6 and 7).

## Results

### Exposure to subclinical doses of epirubicin persistently compromises hUVEC proliferation and viability

EC are essential for organ homeostasis and tissue function, performing localized regulatory roles that maintain adequate exchange of nutrients, gases and signals. hUVEC were used to investigate how this cell type responds to the effects of circulating epirubicin. Time of exposure and concentrations for treatments *in vitro* were determined based on compound half-life and the concentration following dilution of typical clinical doses in the blood stream (Robert 1994), to investigate the effects of the drug at organs distant from the site of infusion. The average concentration found in circulation of patients (10 μg/mL) receiving the typical clinical epirubicin dose (90 mg/kg) proved too toxic for a hUVEC monolayer, and most cells did not survive the treatment (data not shown). Therefore, confluent monolayers were treated for 30 min with lower doses of epirubicin (3 μg/mL or 5 μg/mL), which was subsequently replaced with drug-free growth medium.

Total cell counts following treatment was shown to decrease after 1 d for cells treated with 5 μg/mL (**Figure 1A**). This is also illustrated by a gradual decrease in confluency (**Supplementary Figure S1A**), where more space is observed between cells. For the first 2 d of recovery from epirubicin exposure, attached cells remained viable (as determined by Trypan blue exclusion method), but viability decreased in treated cells after 3 d (**Figure 1B**), suggesting that the cytotoxic effects of the drug on hUVEC can persist for several days following exposure. TUNEL staining of hUVEC following epirubicin treatment shows, predictably, a dose-dependent increase in hUVEC apoptotic cell death (**Figure 1C);** cell number per field of view, quantified by batch analyses (ImageJ) (**Figure 1D**, upper panel) decreases with increased epirubicin dose, with a larger proportion of dead or dying cells particularly significant in cells treated with 5 μg/mL epirubicin. Cells that persist following 3 μg/mL show proportionally less TUNEL staining and appear to remain viable for longer (**Figure 1D**, lower panel).

**Figure 1.**
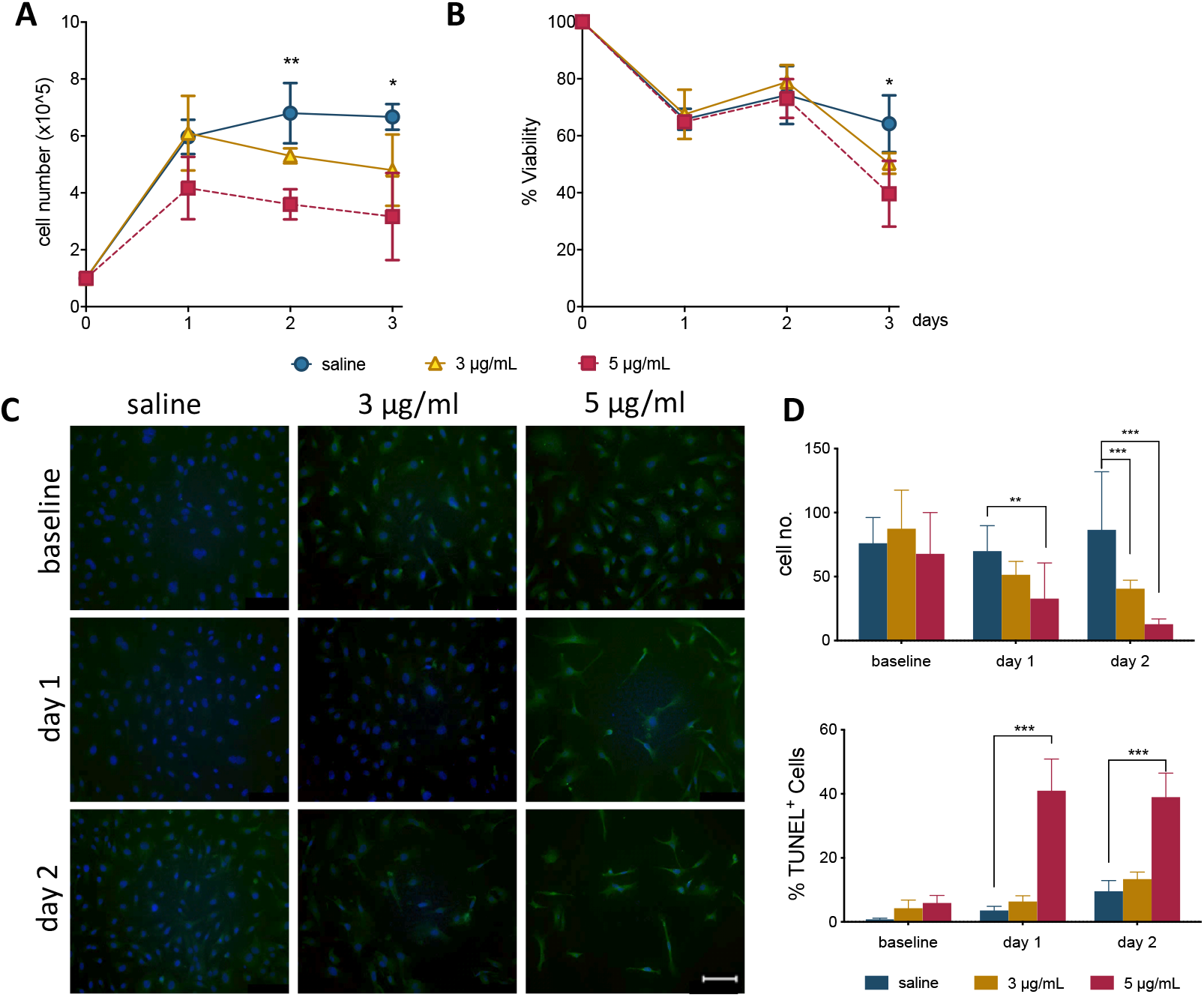
Epirubicin treatment reduces hUVEC proliferation and viability. Confluent hUVEC monolayers were exposed for 30 min to epirubicin (3 or 5 μg/mL) or equal volume of saline. **(A)** Total number of attached cells were counted immediately after treatment (baseline) and again following 1 d, 2 d or 3 d of recovery in drug-free medium. **(B)** Cell viability was assessed using Trypan Blue exclusion. Data displayed as mean ± SD, significance assessed by t-tests and * p < 0.05, ** p < 0.01, n=3. For panels (A) and (B), no significant difference was observed between saline and 3 μg/mL, asterisks represent significance between saline and 5 μg/mL epirubicin. **(C)** Representative images of TUNEL-stained control and epirubicin-treated hUVEC, scale bar = 100 μm. **(D)** Number of cells per field of view (upper panel) and relative proportion of TUNEL positive cells detected per field of view, represented as average percentage (lower panel). Image analyses and quantification done using ImageJ (8.0). Statistical significance was assessed by Two-Way ANOVA with Bonferroni Correction, and *p < 0.05, **p < 0.01, ***p < 0.001.

### Viable EC exposed to epirubicin become senescent

During recovery following exposure to epirubicin, changes in confluency and cell morphology were observed, where cells appear enlarged and flattened (**Supplementary Figure S1B**), usually characteristic of cells undergoing senescence. It has been shown in other models that chemotherapy may cause senescence in somatic cells (Ewald, Desotelle et al. 2010), and this was investigated as a possible effect on hUVEC. The well-established Senescence-Associated β-Galactosidase (SA-β-Gal) method was used to detect senescing hUVEC following epirubicin treatment (**Figure 2A**). Total number of cells per field of view were quantified (**Figure 2B**, upper panel), and the relative proportion of senescing cells (**Figure 2B**, lower panel) is seen to dramatically increase immediately after exposure and is maintained for at least 2 d after drug removal (**Figure 2B**). These results show low concentrations seen in small microvascular networks distant from injection sites can trigger endothelial senescence. Cell senescence is associated with a secretory pattern that is primarily pro-inflammatory (senescence associated secretory profile, or SASP, (Coppé, Patil et al. 2008, Kuilman, Michaloglou et al. 2008, Wajapeyee, Serra et al. 2008)). An angiocrine screen (**Figure 2C**) was performed on hUVEC-conditioned medium, obtained from cells that recovered for 1 d in drug-free medium to assess EC-derived signaling, and shows that exposure to 3 μg/mL of epirubicin caused a significant increase in the secretion of many pro-inflammatory cytokines (**Figure 2D**), consistent with SASP signature, and including monocyte/macrophage recruitment and activation factors such as CCL2 and granulocyte colony stimulating factor (G-CSF) (Coppé, Patil et al. 2008, Eggert, Wolter et al. 2016), as well as pro-inflammatory cytokines IL-6 and IL-8.

**Figure 2.**
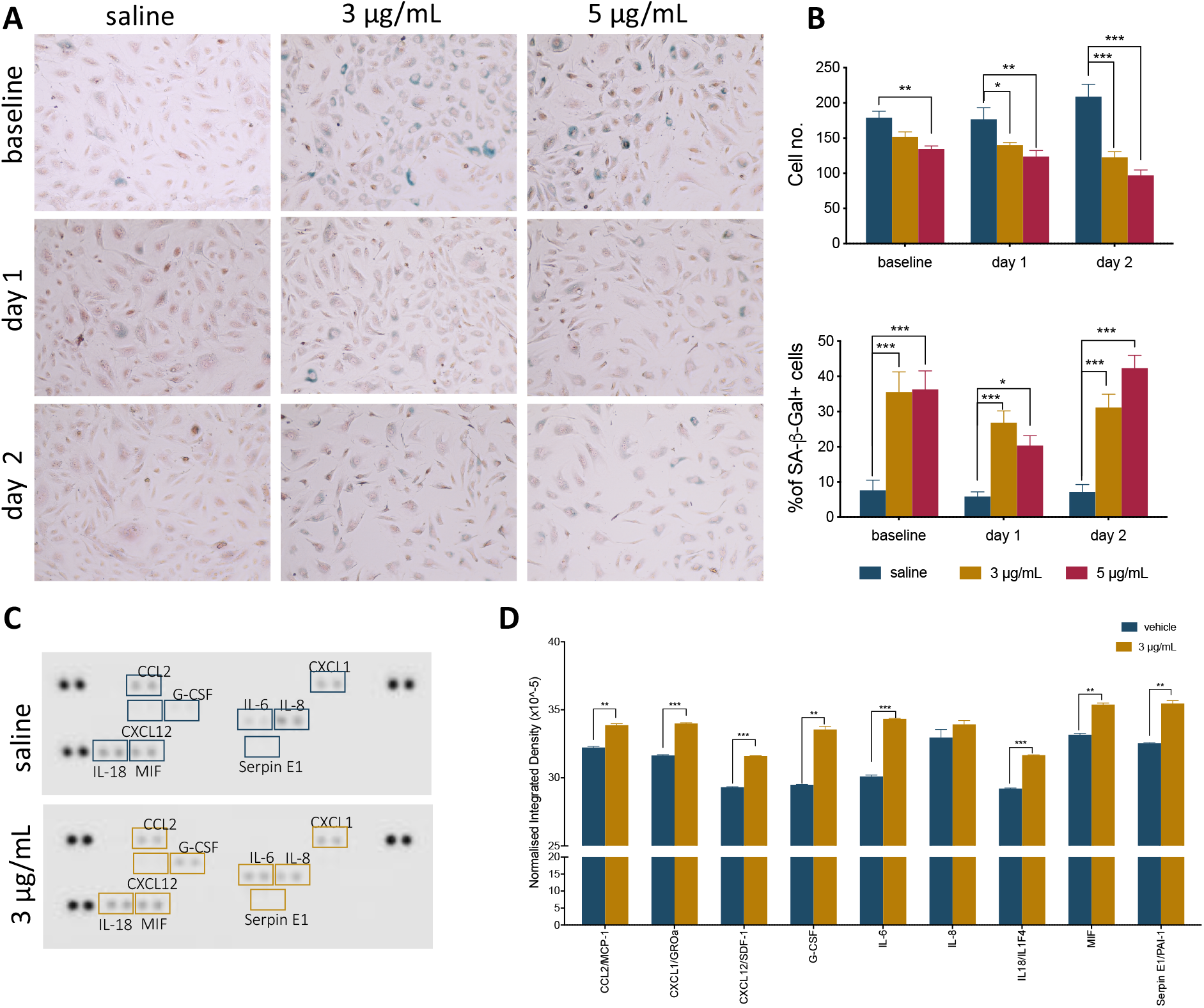
Epirubicin treatment induces senescence in hUVEC. **(A)** Representative images of saline and epirubicin-treated hUVEC detected using senescence marker SA-β-gal, and counter-stained with Nuclear Fast Red. Images were obtained on an EVOS brightfield microscope at 20 × magnification. **(B)** Number of cells per field of view (upper panel) and the average percentage of SA-β-gal positive cells (lower panel); Data displayed as mean ± SEM (n=3). Statistical significance was assessed by Two-Way ANOVA with Bonferroni Correction, and *p < 0.05, **p < 0.01, ***p < 0.001. **(C)** Blot image of cytokine panel of hUVEC-conditioned medium collected 1 d after 30 min treatment with saline or 3 μg/mL epirubicin. **(D)** Densitometry analysis of detectable targets in (C) was carried out using the raw integrated density function on ImageJ software. Data displayed as mean ± SD. Unpaired t-tests were used for analysis, and *p < 0.05, ** p < 0.01, n = 2.

### Exposure to epirubicin impairs endothelial function

To further evaluate if the effects of epirubicin influences endothelial function, essential parameters of endothelial behavior were measured. All functional parameters were performed in cells for which viability was confirmed (**Supplementary Figure S2A**).

The ability to form and regenerate monolayers is essential in maintaining vascular integrity during angiogenesis and wound healing, and a property that relies on endothelial migratory capacity. This parameter was quantified using a scratch assay, in the presence of mitomycin, and wound closure was measured over time (**Figure 3A**). Cells exposed to epirubicin showed a delay in wound closure, but significance was only established for cells treated with a higher dose (5 μg/mL), when the assay was performed after 1 d of recovery (representative images for this time point are shown in **Figure 3B**). Barrier function was also assessed following 1 d recovery, using an assay in which the movement of FITC-labelled 70 kDa Dextran across a hUVEC monolayer was measured in real-time. Results presented in **Figure 3C** show that permeability increased significantly following epirubicin treatment. This phenotype is associates with internalization of VE-cadherin, the key adherens molecule in EC, responsible for vascular integrity and stability of endothelial cell-cell adhesion. VE-cadherin signal shifts from the membrane to cytoplasm following epirubicin treatment, and this correlated with cell dissociation, observed by the appearance of gaps between hUVEC. (**Supplementary Figure S2B)** Angiogenic potential was also assessed using a tube formation assay (**Figure 3D**), where cells were assayed immediately after treatment (upper panel), and after 1d (middle panel) and 2 d (lower panel) recovery. Images were collected at different time points and analyzed using ImageJ software to systematically quantify number of branches (left) and junctions (right) formed in each condition. Tube formation in saline controls peaked between 3 h and 5 h, thus representative images for the 5 h time point are shown (**Figure 3E**), which also represents the time point at which most significant differences were seen across experimental groups. hUVEC ability to form networks is impaired immediately and significantly after treatment (**Figure 3D**, upper panel, baseline), where untreated hUVEC form more complex patterns with a higher number of branches and junctions. Interestingly, following recovery for 1 d (**Figure 3D**, middle panel), cells exposed to 5 μg/mL of epirubicin show a higher number of branches than those exposed to 3 μg/mL. After 2 d in drug-free medium, cells treated with the lower epirubicin dose show an angiogenic potential comparable to that of untreated cells (**Figure 3D**, lower panel), whereas cells exposed to 5 μg/mL lose the ability to form networks. Representative figures for each experiment (baseline and two recovery time points) are shown in **Figure 3E**.

**Figure 3.**
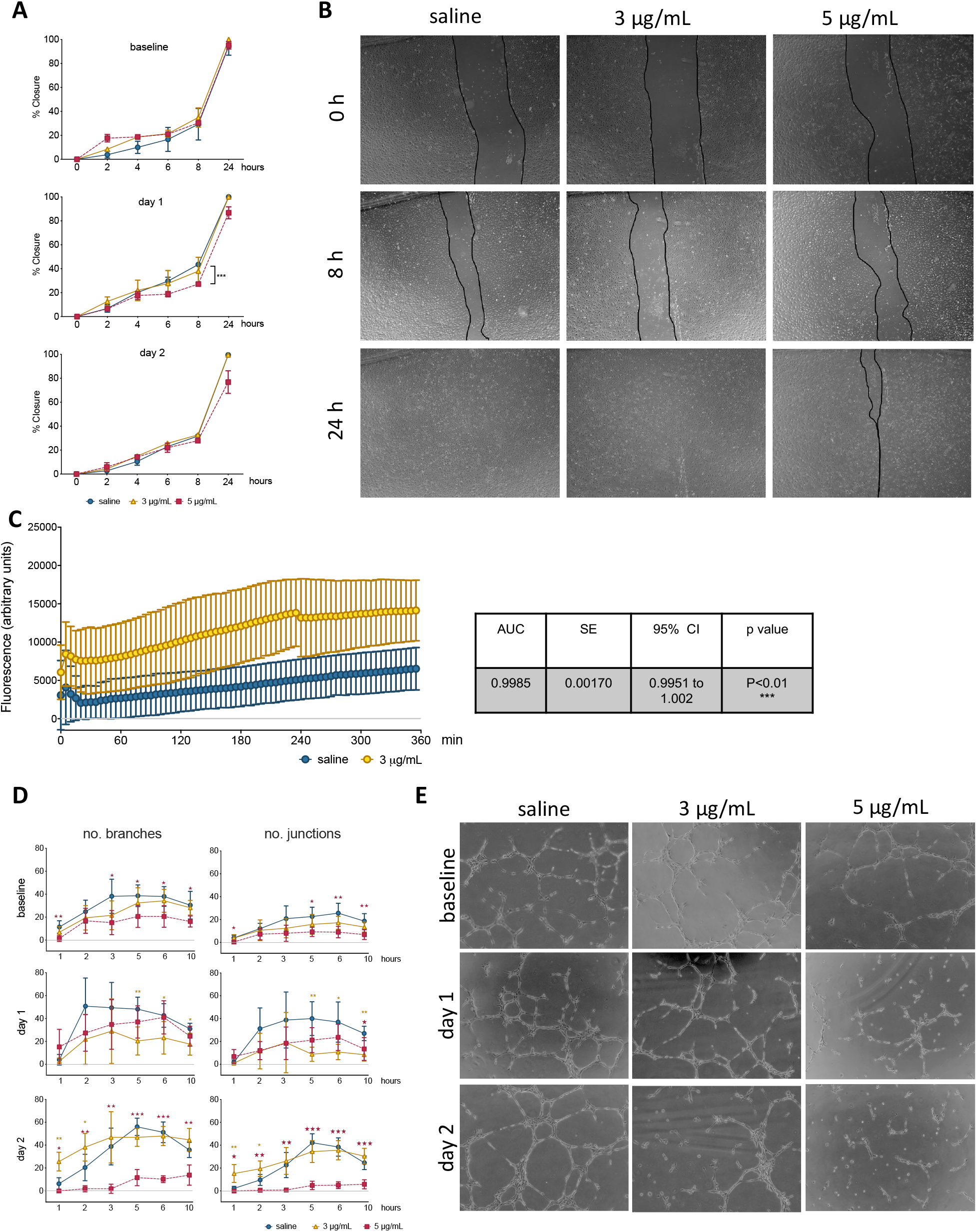
Low dose epirubicin compromises endothelial migration, barrier function and angiogenic potential. **(A)** Migration assay with viable epirubicin or saline-treated hUVEC, performed in the presence of Mitomycin-C. Assays were performed immediately after exposure (baseline, top panel), 1 d (middle panel) or 2 d (lower panel) recovery in drug-free medium. Images of 2 wells per treatment were collected at regular intervals for up to 24 h, and the area of the scratch was quantified using ImageJ software and converted to percentage of closure. Data displayed as mean % closure ± SD; experimental groups were compared at each time point using unpaired t-tests, and *p < 0.05, **p < 0.01, ***p < 0.001. **(B)** Representative images of migration assay from cells 1 d after treatment. **(C)** Permeability assay detecting FITC-dextran (70 kDa) signal as the compound moved through hUVEC monolayer 1 d after epirubicin treatment. Area under the curve (AUC) was compared between treatments, and standard error of the mean (SE) were calculated using GraphPad Prism and shown in adjacent table. **(D)** Tube formation assays were performed immediately after incubation (baseline), 1 d and 2 d after recovery in drug-free medium. Number of branches and junctions were quantified using the ‘Angiogenesis Analyser’ plug-in for ImageJ; Data represents mean ± SD (n=5). Red stars represent comparisons between saline and 5 μg/mL epirubicin, yellow asterisks represent comparisons between saline and 3 μg/mL epirubicin. Unpaired t-tests were used for analysis of each time point, *p < 0.05, **p < 0.01, ***p < 0.001. **(E)** Representative images of tube formation assays at the 5 h time point.

In order to investigate if these functional differences were seen to have an underlying transcriptional control, a Transcription Factor (TF) activation assay was carried out for cells treated with the lower dose of 3 μg/mL epirubicin, following 1 d recovery (**Supplementary Figure S3A**). Most TF activity decreases in response to epirubicin, with the most striking decrease observed for SMAD and Sp1, both of which have been associated with VEGF activity (Pagès and Pouysségur 2005, Lin, Xie et al. 2016). Consistently, 1 d after treatment, VEGF mRNA levels decrease dramatically. However, after 2 d an increase is observed to much higher levels than saline controls, and more so in cells treated with the lower epirubicin dose (3 μg/mL). One other key molecular marker of EC health and function is the endothelial nitric oxide synthase (eNOS), which is essential for vascular tone, intercellular signaling and local perfusion homeostasis (Heiss, Rodriguez-Mateos et al. 2015); eNOS transcript levels decrease in a dose-dependent manner, but also transiently (with a slight increase at day 2 compared to day 1) (**Supplementary Figure S3B**).

In summary, these data suggest that there is both a dose- and time-dependent effect on endothelial response to epirubicin. Cells exposed to higher concentrations respond to angiogenic stimuli during the early recovery phase, but fail to maintain that ability (**Figure 3D**, middle panel). However, cells that are exposed to milder toxicity (3 μg/mL) resume the angiogenic potential after 2 d to levels comparable to the saline-treated controls, and this is consistent with a decrease (day 1) followed by an increase (day 2) in a major angiogenesis mediator (VEGF, **Supplementary Figure S3B**).

### Metabolic activity is compromised in hUVEC exposed to epirubicin

It has been demonstrated that endothelial metabolism governs, to a significant extent, endothelial function (Bierhansl, Conradi et al. 2017). To evaluate metabolic activity in hUVEC exposed to epirubicin, cells were subjected to mitochondrial stress tests (**Figure 4A, B**) and glucose stress tests (**Figure 4C, D**). These parameters were evaluated immediately after treatment (baseline) or after cells had been allowed 1 d to recover, using a Seahorse metabolic analyzer (Agilent).

**Figure 4:**
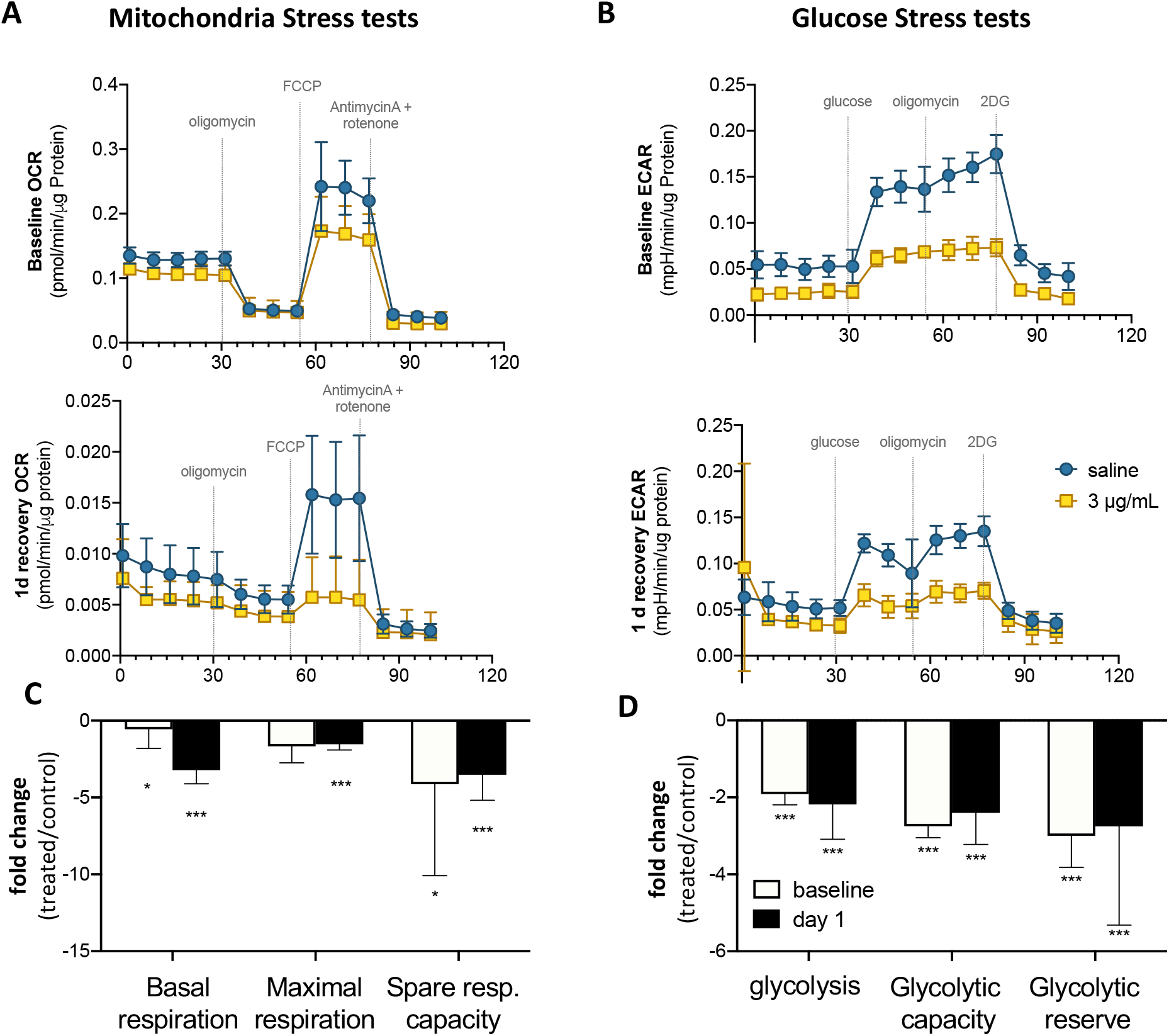
Endothelial metabolism is significantly reduced following epirubicin exposure. hUVEC treated with saline or epirubicin for 30 min were analyzed immediately (baseline) or after 1 d in drug-free medium. **(A)** Mitochondria stress tests performed at baseline (upper panel) or after recovery (lower panel). Data presented as mean ± SD, n = 8 wells. **(B)** Summary chart showing mean fold-change ± SEM of mitochondrial respiration parameters (treated/control) **(C)** ECAR measurements obtained from glycolysis stress test are shown for baseline and following 1 d recovery, (upper and lower charts, respectively); as in (A), data represents mean ± SD, n = 8 wells. **(D)** summary graph of relevant glycolytic parameters, showing mean fold-change (treated/control) ± SEM. Student’s t-tests were used to assert significance of differences between treatment and saline controls, ***p < 0.0001 and *p < 0.05.

In both time points (baseline and 1 d recovery), cells exposed to epirubicin show a significant decrease in oxygen consumption rate (OCR) (**Figure 4A**), but the decrease in mitochondrial respiratory parameters compared to control cells is more marked 1 d after treatment (**Figure 4B**), suggesting that mitochondrial reprogramming continues several hours after exposure in viable EC. Glycolytic activity is also seen to decrease in response to epirubicin (**Figure 4C**), however, unlike the gradual changes seen in oxidative phosphorylation, this shift happens sharply and immediately after the treatment (baseline) and is maintained for at least 1 d after exposure (**Figure 4D**). In both cases, there is a clear decrease in metabolic activity, which can be a result of decreased energetic demand following onset of senescence, or, conversely, a senescent phenotype as a result of a decrease in energy availability. Consequent decrease in ATP availability is consistent with a significantly reduced (energy demanding) transcriptional activity (**Supplementary Figure S3**).

### Lung MVEC isolated from mice treated with epirubicin show decreased viability and migration

EC are a very heterogeneous population, and MVEC are more plastic and versatile than their venous counterparts (Dejana, Hirschi et al. 2017); these cells are, however, the ones that locally refine and coordinate tissue demands with systemic availability and distant signals. To assert if the cytotoxic insult is similarly perceived and responded to by MVECs *in vivo*, naïve C57BL/6 mice were treated with a low clinical dose of epirubicin (approximately at 13.6 mg/kg). Lung MVECs were subsequently isolated either 1 d or 7 d after injection (**Figure 5A**), to assess immediate and persistent effects. Following the establishment of a MVEC culture, cells were trypsinized and plated at the same density (passage 1, P1), and total and viable cells were counted every day for 3 d (**Figure 5B**). Number and viability of cells isolated 1 d after treatment are similar to those obtained from mice that received saline injections, but interestingly, lung MVEC obtained from mice one week after epirubicin injection proliferated slower and had slightly compromised viability, which is significant after 3 d in P1.

**Figure 5.**
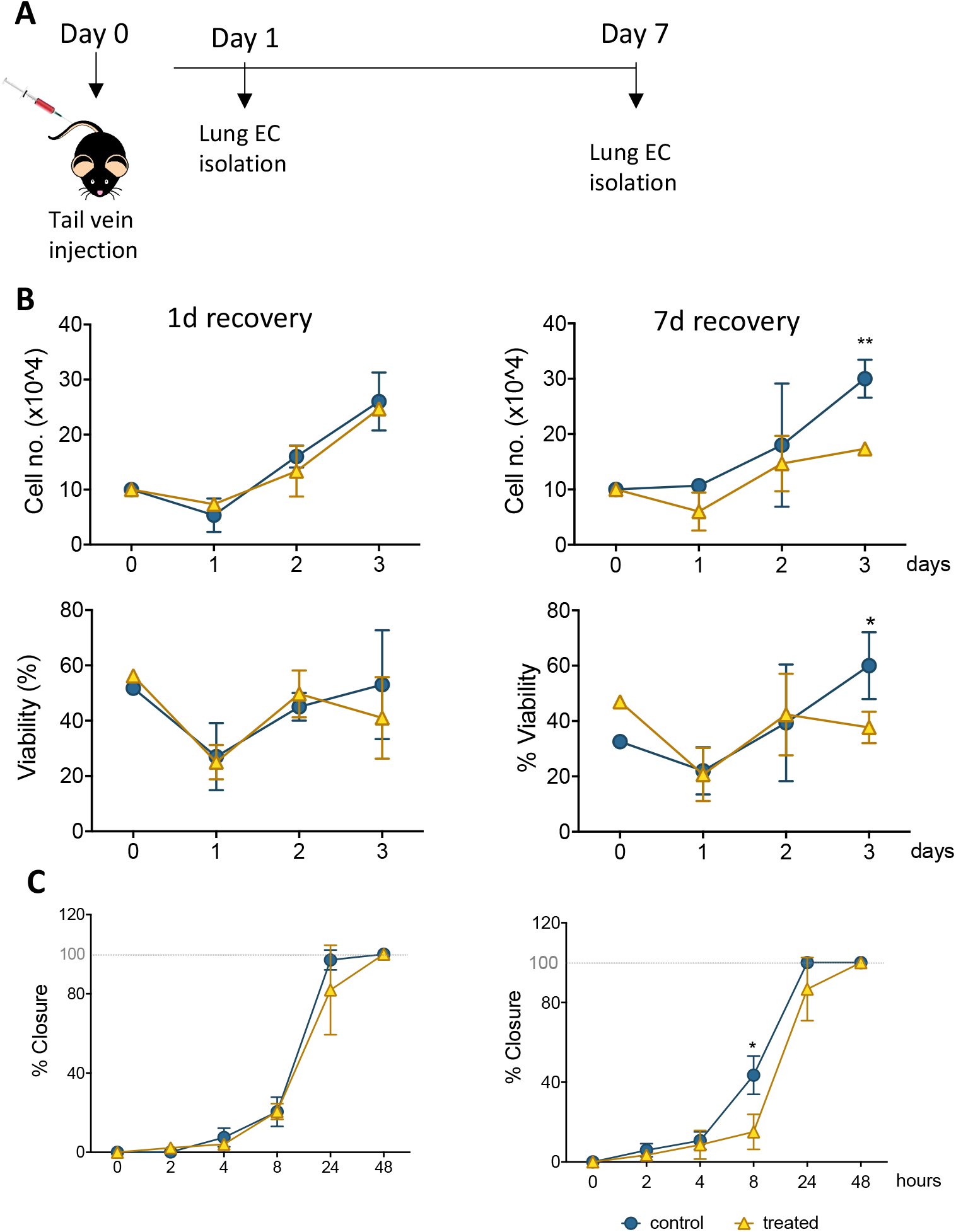
Primary lung MVEC isolated epirubicin-treated mice: effects on viability and migration. **(A)** Experiment outline EC exposure *in vivo;* C57BL/6 mice received intravenous epirubicin or saline control, and lung MVEC were isolated 1 d and 7 d post injection. Lung MVEC cultures were established and assays performed with cells at P1. **(B)** Cells isolated 1 d (left) and 7 d (right) were used to quantitatively compare their proliferation (upper panels) and viability (lower panels). **(C)** Migration assay of P1 lung MVEC obtained after 1 d or 7 d of mice that received saline control or epirubicin injections. Data displayed as mean ± SD of 3 independent replicates, unpaired t-tests used for analysis, * p<0.05, ** p<0.01.

To assess if this change in proliferation had functional implications, a migration assay was performed with P1 primary lung MVEC, in the presence of mitomycin C (as in **Figure 3**). Again, as seen for proliferation and viability, cells isolated from mice 7 d after epirubicin injection had significantly impaired migration compared to the ones obtained from saline controls (**Figure 5C**). These data demonstrated that changes in MVEC behavior occur *in vivo*, but are not seen with immediate effect, suggesting that adaptation involves endothelial reprogramming and not just a transient response. Interestingly, the effect on cells isolated 1 d after injection is not seen even after 1 week in culture, and it is possible that the adaptive response seen *in vivo* is interrupted or reversed by excess growth factors or nutrients present in growth medium.

### Epirubicin causes a decrease in lung vascular density

The lung microvasculature is at the heart of lung function, and efficient gas exchange relies entirely on the seamless interface between a large epithelial surface in contact with a correspondingly large endothelial surface, accomplished with a dense endothelial network. Whole lung tissue from saline control and epirubicin-treated mice were fixed and paraffin-embedded for immunofluorescent analysis. Two groups of mice received one injection of epirubicin (as in **Figure 5**) and tissue was collected after 1 d or 7 d. A third group of mice was treated with two epirubicin doses 3 d apart, to evaluate cumulative effects (**Figure 6A**). Sections of 10 μm were stained for EC (Podocalyxin, (Li, Li et al. 2001) and pericytes (α-SMA, (Babai, Musevi-Aghdam et al. 1990); nuclei were detected with DAPI and representative images are shown in **Figure 6B**. Images were systematically analyzed in batches to quantify multiple vascular parameters (**Figure 6C**), including vascular density (left panel), vessel diameter (middle panel) and pericyte:EC ratio, as a measure of pericyte coverage and vascular integrity. Consistent with what was observed in the *ex vivo* experiments (**Figure 5**), a significant decrease in vascular density (**Figure 6C**, left) was seen only after 7 d, but not immediately after injection (1 d). Surprisingly, animals that received two epirubicin injections within the same week, also had reduced vascular density but not different from mice that received only one treatment. This may suggest that cells are either less responsive, or that indeed it takes more than 3 d for a cumulative effect to manifest *in vivo*.

Vessel diameter (**Figure 6C**, middle) was not altered, and pericyte:EC ratio (**Figure 6C**, right) was also comparable between all treatment groups, which suggests a vessel pruning effect (reduction in number of vessels) and not a change in integrity or caliber of blood vessels present.

**Figure 6.**
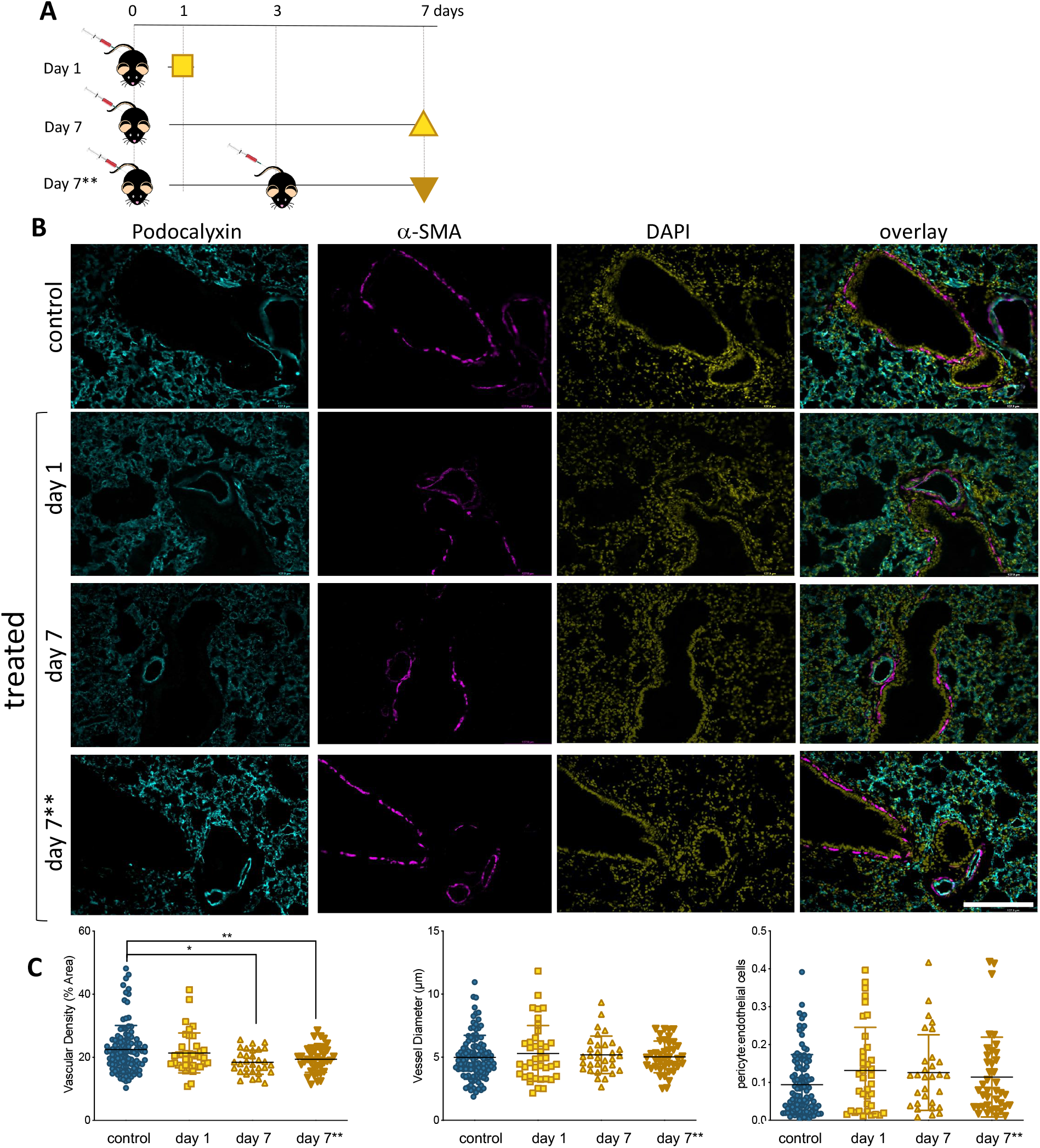
Lung vascularity decreases in epirubicin-treated mice 7 days post injection. **(A)** Experiment outline: 6-8 week-old C57BL/6 mice received intravenous epirubicin or saline injections, and tissue harvested and analyzed 1 d (day 1) or 7 d (day 7) post injection. A third group of animals were re-injected at day 3 and lungs harvested on day 7 (day 7**). **(B)** Representative images of lung sections stained with endothelial marker (Podocalyxin, Cyan), pericyte marker (α-SMA, Magenta) and counterstained with nuclear marker DAPI (Yellow); Scale bar = 200 μm **(C)** Vascular parameters were analyzed using Image J. Vascular density (average % area ± SD), vessel diameter (average μm ± SD) and Pericyte coverage (displayed as a ratio of pericyte: EC signal). Charts combine data from 8 sections per animal, 3 images per section (4-5 animals per group, saline controls are combined); statistical significance was assessed by one-way ANOVA with Dunnett’s Multiple Comparison Test, and *p < 0.05, ** p < 0.01.

### Epirubicin treatment promotes myeloid cell infiltration into the lung

Our results in hUVEC show that one of the most striking reprograming events taking place in response to sub-lethal epirubicin exposure is that of driving senescence. Consistent with a senescent-like morphology and positive staining for senescence marker SA-β-gal (**Figure 2**, **Supplementary Figure 1**), a pro-inflammatory SASP-like cytokine profile was also observed. Using sections obtained from the mice in the previous experimental set-up (**Figure 6A**), we used a general myeloid cell marker (Mac 2, (Ho and Springer 1982)) to stain inflammatory cell infiltration into lungs following epirubicin treatment (**Figure 7A**). The number of after analyses and + cells were systematically quantified using batch analyses and ‘particle analysis’ function (ImageJ). Unlike what is seen for vascular parameters, myeloid cell infiltration occurs as soon as 1 d following epirubicin treatment, and is maintained consistently higher than the controls after 7 d. Animals in group receiving two injections (7**) also had a higher number of Mac-2+ cells, but less than either of the other two treatment groups. This was unexpected but could be a result of epirubicin treatment compromising survival and/or proliferation of Mac-2+ cells that infiltrate the lung after the first injection. Combined, and correlating with an increase in EC-derived pro-inflammatory cytokine profile (**Figure 2C, D**), these results show that there is an increase in myeloid cell recruitment to lungs of epirubicin-treated animals.

**Figure 7.**
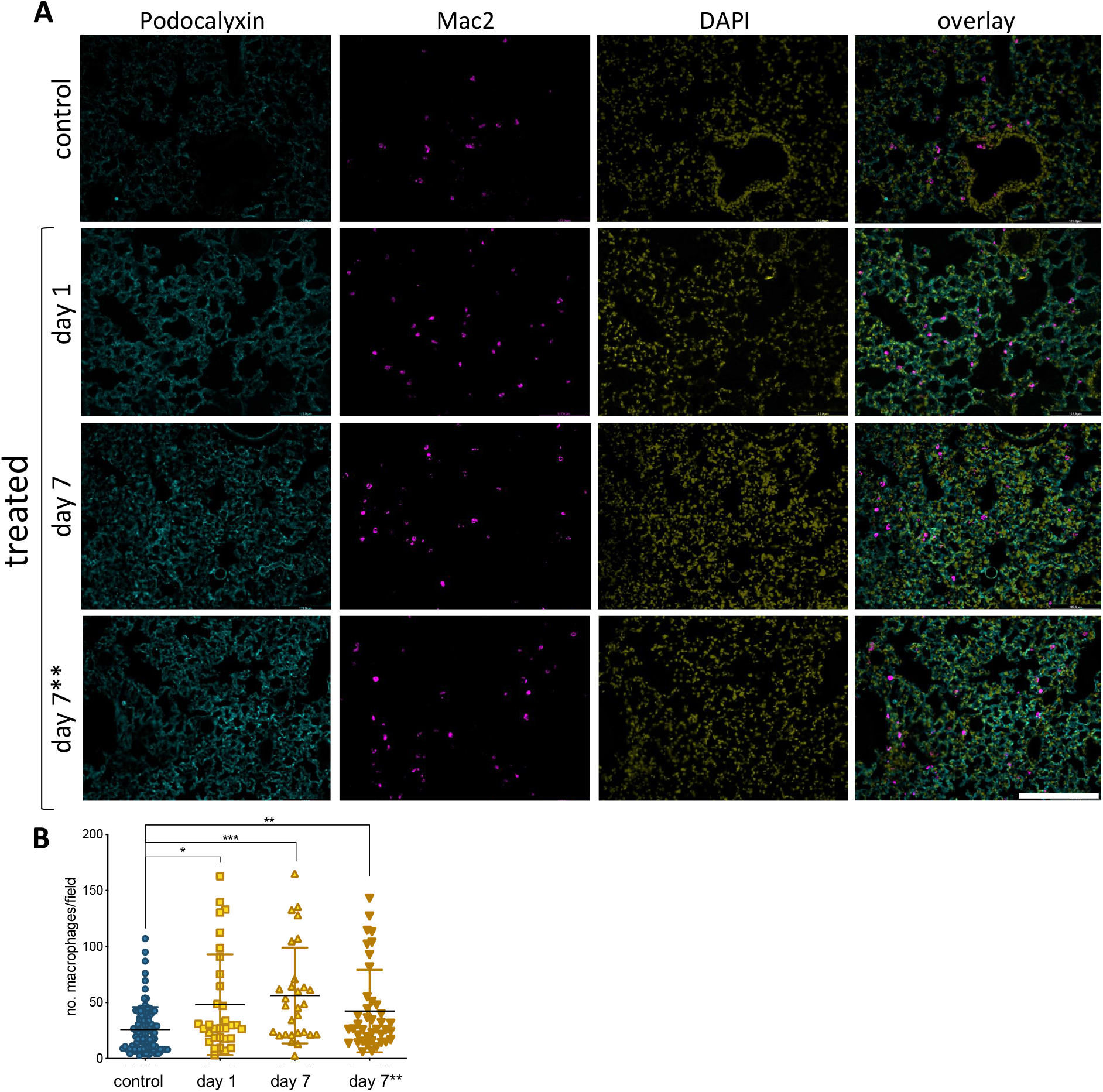
Epirubicin affects the immune cell response in murine lung tissue. **(A)** Lungs sections were stained with Podocalyxin (Cyan), Mac-2 (Magenta) and DAPI (Yellow) and representative images are shown; Scale bar = 200 μm. **(B)** Number of infiltrating myeloid cells was assessed by quantifying Mac-2-positive cells in lung tissue, using Image J software. Charts combine data from 8 sections per animal, 3 images per section (4-5 animals per group, saline controls are combined); Data shown as mean ± SD. Statistical significance assessed by one-way ANOVA with Dunnett’s Multiple Comparison Test, and * p < 0.05, ** p < 0.01, *** p < 0.001.

## Discussion

The vascular endothelium is a critical barrier maintaining homeostasis between circulating blood and surrounding tissues. It is responsible for various functions including maintaining vascular tone, preventing platelet aggregation, and regulating extravasation and angiogenesis, as well as inflammatory responses (Sabbatinelli, Vignini et al. 2017, Kruger-Genge, Blocki et al. 2019), ultimately regulating organ perfusion. The association of anthracycline treatment with endothelial dysfunction at the injection site has been demonstrated (Yamada, Egashira et al. 2012, Sato, Kondo et al. 2014). This study investigates the effects of low dose epirubicin on EC and aimed to model the exposure of distant microvascular networks to this drug. This is important to assess the nature, extent and potential reversibility of vascular responses to epirubicin, and to infer possible secondary disease following cancer treatment, downstream of endothelial reprograming.

As expected, our results show a significant decrease in EC number and viability over time as a result of epirubicin treatment (**Figure 1**): clinical doses of epirubicin can cause significant levels of cell death in hUVEC, but lower doses (3 μg/mL) result preferentially in the induction of senescence (**Figure 2**). This response has been previously associated with treatment with Doxorubicin, and shown to impact on regenerative function (Yasuda, Park et al. 2010, Wojcik, Buczek et al. 2015, Cappetta, Rossi et al. 2018).

Although EC function has been demonstrated to be regulated downstream of metabolic reprogramming (Bierhansl, Conradi et al. 2017), metabolic activity of senescent EC is not particularly well described. Concomitant with a decrease in parameters of EC function assessed either *ex vivo* (**Figure 3**) or in primary cells isolated following *in vivo* exposure (**Figure 5**), we see a dramatic decrease in metabolic activity in hUVEC exposed to 3 μg/mL epirubicin (**Figure 4**). Previous reports have associated senescence with increased metabolic activity due to acquisition of SASP (Wiley and Campisi 2016). Glycolysis was shown to be elevated in senescent human diploid fibroblasts (Bittles and Harper 1984, James, Michalek et al. 2015), however glycolysis upregulation is not typical in senescent EC (Unterluggauer, Mazurek et al. 2008), and instead a decrease in glycolysis has been observed as a result of PFKFB3 activity (Kuosmanen, Sihvola et al. 2018). Here, we show a striking and immediate decrease in glycolytic activity, capacity and reserve (**Figure 4C, D**) that persists for at least 24 h after recovery in complete medium. Mitochondrial respiration also decreases, mostly the basal respiration rates, and at a more gradual pace, becoming significantly more pronounced 1 d after treatment.

The resulting and expectedly sharp drop in ATP availability would result in a decrease in energy-demanding aspects of EC behavior. In fact, a decrease in angiogenic potential and wound-healing ability is seen as a result of epirubicin treatment (**Figure 3** and **Figure 5**), as well as severe downregulation of transcriptional activity (Park, Kim et al. 2013, Zecchin, Kalucka et al. 2017) (**Supplemental Figure S3A**). The most remarkable decrease is shown in the TF activity of SMAD and Sp1, both of which linked to vital EC functions. SMAD is a key regulator of angiogenesis (Lin, Xie et al. 2016), vascular stability and vascular remodeling (Lan, Liu et al. 2007, Itoh, Itoh et al. 2012); importantly for patients undergoing treatment for cancer, SMAD deletion has been associated with increased metastatic disease (Yang, Wang et al. 2017).

Sp1 has been associated with cell proliferation, differentiation, apoptosis and senescence (Beishline and Azizkhan-Clifford 2015), and is also an important transcriptional regulator of VEGF, and thus plays a direct role in angiogenesis (Ko and Kim 2018). It may indeed play also an indirect role in management of cellular energy metabolism, as decreased VEGF levels (as seen in **Supplemental Figure S3**) have been shown to result in decreased glycolysis (De Bock, Georgiadou et al. 2013). VEGF inhibition has also been associated with endothelial senescence, via elevated intracellular reactive oxygen species (ROS) (Mongiardi, Radice et al. 2019).

Importantly, cell senescence is associated with an inflammatory microenvironment (Coppé, Patil et al. 2008, Kuilman, Michaloglou et al. 2008, Wajapeyee, Serra et al. 2008), identified in epirubicin-treated hUVEC conditioned medium (**Figure 2C, D**). The effects of inflammation in vascular function are notably reciprocal, and associated with a myriad of chronic health conditions, such as diabetes, ageing and neurodegeneration and multiple respiratory syndromes (Augustin and Koh 2017, Reiterer and Branco 2020).

Epirubicin, although with much milder associated cardiotoxicity than other anthracyclines, is shown here to promote endothelial secretion of IL-6 and IL-8 (Childs, Durik et al. 2015), previously linked to senescence and increased tumorigenesis (Ortiz-Montero, Londoño-Vallejo et al. 2017), which in turn may be associated with increased risk of metastatic disease. Additionally, monocyte recruitment and activation signals, such as CCL2 and G-CSF (Coppé, Patil et al. 2008, Eggert, Wolter et al. 2016), (**Figure 2**), are also pro-tumorigenic. These increased monocyte recruitment signals seen in hUVEC treated with epirubicin correlate with significantly elevated myeloid cell infiltration (Mac-2+ cells) into lungs of epirubicin-treated mice compared to saline controls (**Figure 7**). This effect, seen 24 h after injection, is not reversed within one week. Interestingly, however, it is not further increased if the animals receive a second injection 3 d apart. This can be a result of senescent cells being either unresponsive to the subsequent stimulus, or unable to increase the pro-inflammatory signal. It is also possible that myeloid cells present in the tissue following the first injection will undergo apoptosis in the acute phase of response to the second injection. This can only be confirmed with additional time points and co-staining experiments, in which the response of other cells types is evaluated.

Microvascular EC in live tissue are more plastic and versatile than their venous counterparts(Augustin and Koh 2017); also, unlike hUVEC maintained *ex vivo*, these cells are able to rely on the coordinated response of multiple cell types, which we anticipated would have a protective effect on EC viability and function. This is supported by the fact that, on one hand, the effects seen on vascular regression (**Figure 6**) or inflammatory cell infiltration (**Figure 7**) are not intensified with a second injection; however, and very importantly, the effects are also not reversed, which supports the proposition of tissue remodeling and long-term changes in microvascular and organ function following treatment.

One of the major risks associated with cancer is distant recurrence, and recent evidence indicates that chemotherapy treatment can potentiate metastatic disease (Chabner 2018). For example, Paclitaxel and Doxorubicin have been shown to facilitate intravasation in human xenograft models of breast cancer, with concomitant increase in secondary tumors (Karagiannis, Pastoriza et al. 2017). Besides EC, other cells such as macrophages secrete proinflammatory cytokines such as IL-6 and IL-12 following chemotherapy (Bryniarski, Szczepanik et al. 2009), further promoting cancer cell proliferation and metastasis (Liu, Lin et al. 2015). These studies focus on the effects of chemotherapy in the tumor vasculature but overlook that of tumor-free organs. Disrupting MVEC integrity, signaling and barrier function (**Figure 3B** and **Supplementary Figure S2B**) happens at low dose exposure to epirubicin in EC. Little is known about the mechanisms that lead to drug-promoting metastasis via vascular rearrangements, but these results suggest that endothelial disruption by sublethal exposure to epirubicin could facilitate extravasation and metastatic colonization, especially when combined with an inflammatory microenvironment. Further investigation is needed to validate this hypothesis and to determine the association between cytotoxic treatment and disease progression.

Clinically, cancer patients are exposed to long term, cumulative treatment strategies. Our results show that despite removal of initial epirubicin exposure, over time hUVEC do not recover *in vitro*. Furthermore, we observed a stable decline in lung vascular density following treatment *in vivo*. We propose that vascular responses to epirubicin, and possibly other anthracyclines, result in molecular reprogramming and adaptations that are likely to result in tissue remodeling and long-term changes in vascular and organ function, thus underlying secondary pathologies following treatment, and potentially also affecting risk for metastasis. Further understanding of vascular responses in organ microenvironment will assist in prediction and prevention of chemotherapy-associated long-term secondary pathologies and survival-associated morbidities.

## Author Contributions

AJE and TMcE performed cell proliferation and viability assays. AJE performed all functional assays (angiogenesis, migration and permeability), as well as cytokine and transcription factor activation screens; TMcE performed all cytochemistry and hUVEC staining, imaging and quantification (TUNEL, SA_β-Gal, VE-cadherin) and RTqPCR. AJE and GA performed animal experiments and collected tissue for IF. AJE, GA and NT processed and sectioned tissue for IF; AJE and GA isolated and maintained primary EC for viability, proliferation and migration assays. AB designed, optimized and performed all the metabolism assays and analyses. AE performed, imaged and analyzed all murine lung IF staining. CB designed and supervised the study. CB, AJE and TMcE wrote the manuscript. All authors made contributions to the final version and proof-read the manuscript.

## Acknowledgments

The authors would like to thank Niamh McGuckin, Lowri Edwards and Aaron Johnston for their contribution towards optimization of staining protocols and image analyses and quantification, as well as for their thorough preparation of detailed methodology for batch analyses using ImageJ software. We would also like to acknowledge Dr. Moriz Reiterer for his invaluable contribution to the optimization of isolation and maintenance primary MVEC, as well as technical assistance with the seahorse metabolic assays and analyses.

## Supplementary Material

Please refer to Supplementary material for control experiments and data in support of that presented in main figures.

---

**Supplementary Figure S1.**
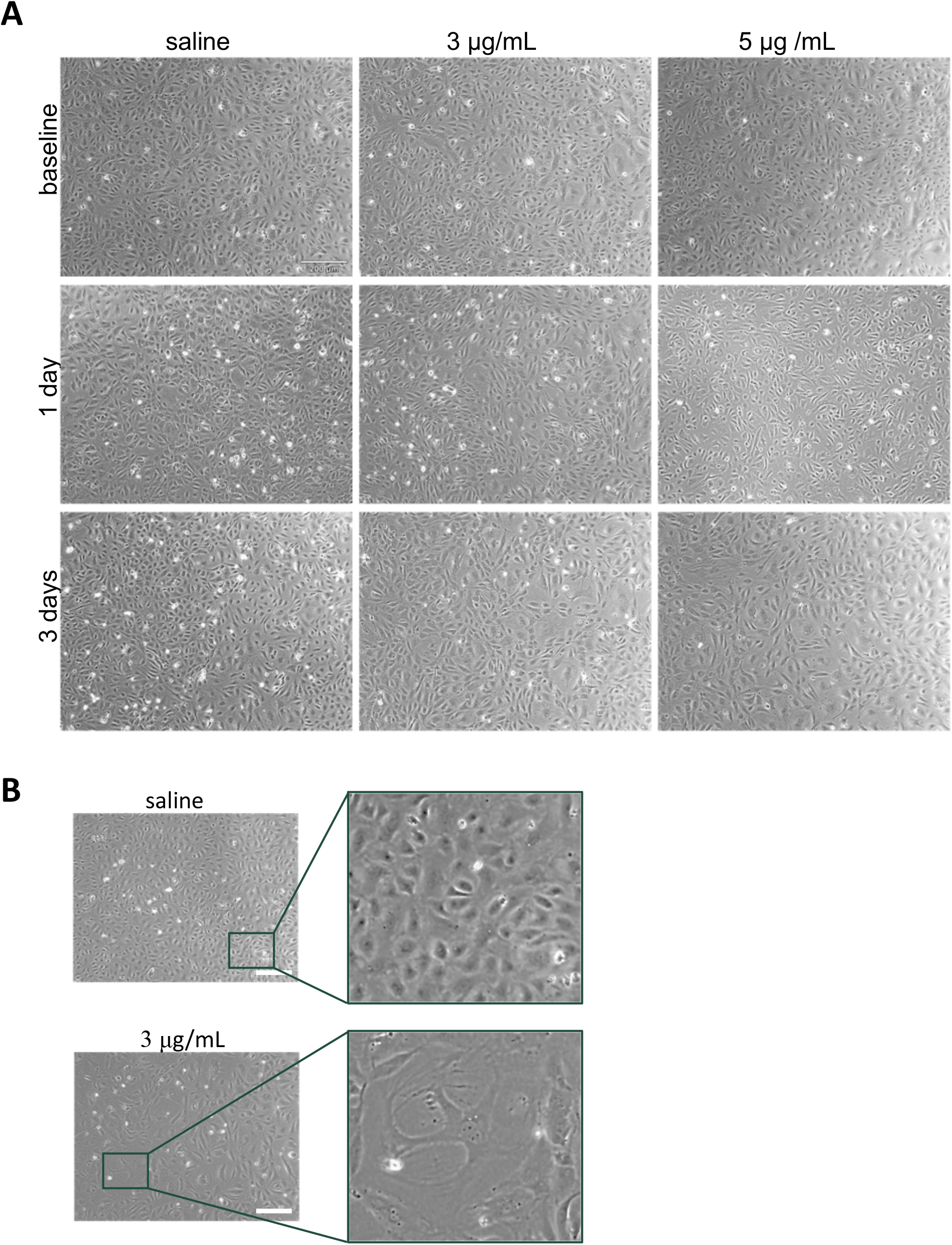
Effects of epirubicin exposure on cell density and morphology. (A) Confluency images of various time points throughout the experiment. Changes in confluency were observed using an EVOS phase contrast microscope (10x magnification). (B) Representative images and enlarged insets to illustrate senescent-like morphology of endothelial cells following exposure to Epirubicin. Images were taken at 10 × magnification 7 d following initial treatment with saline or 5 μg/mL Epirubicin. Scale bar represents 200 μm.

**Supplementary Figure S2.**
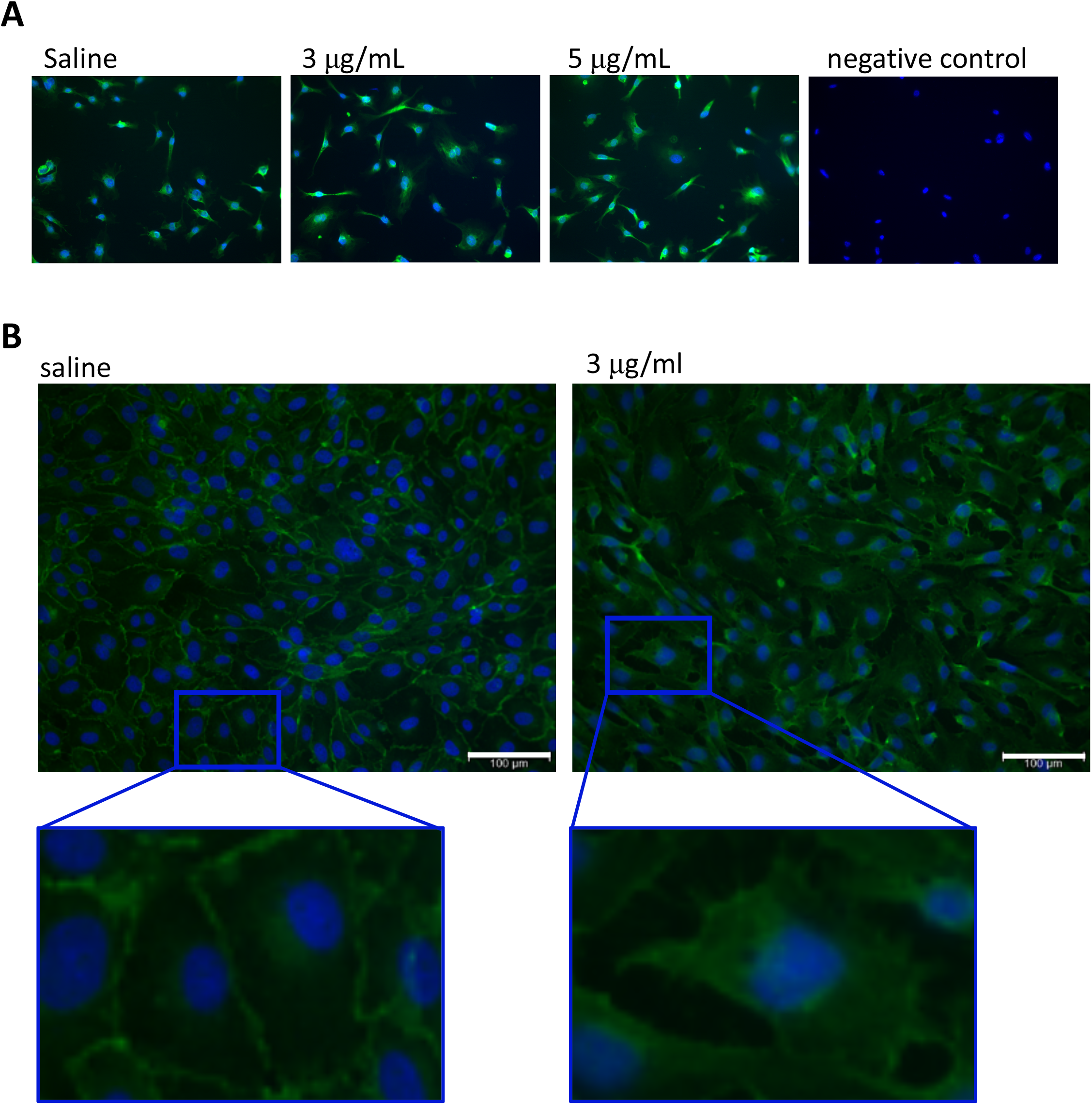
Viability control prior to functional assays (A) and effects of epirubicin on EC junctions (B) (A) Cells used for functional assays (main Figure 3) were stained with Calcein AM at the last time point to ensure viability of experimental cell population throughout the assays. Blue = DAPI, Green = Calcein AM. (B) VE-Cadherin staining of hUVEC 1 d after treatment with 3 μg/mL Epirubicin of respective saline control.

**Supplementary Figure S3.**
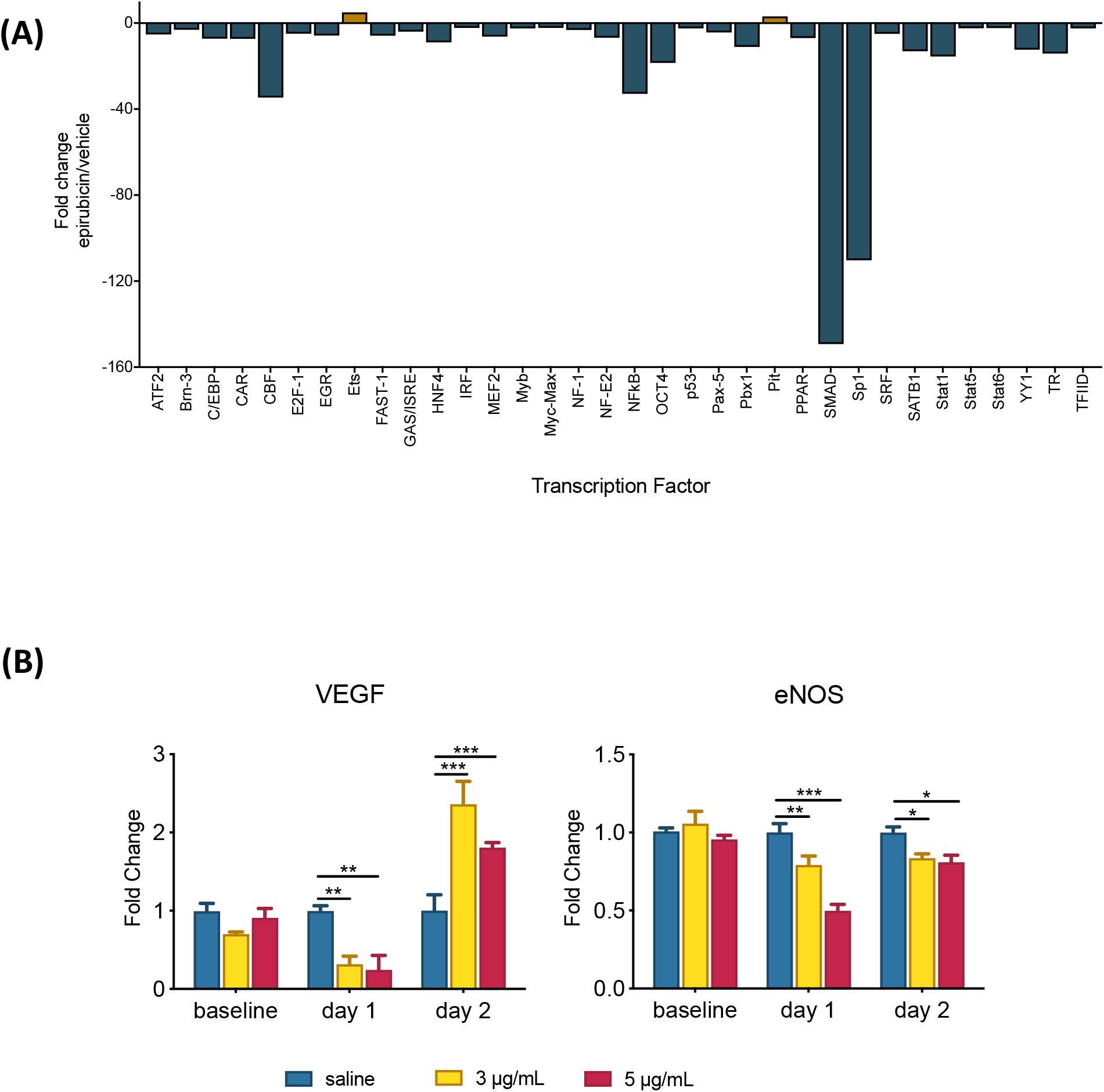
Epirubicin affects transcriptional activity in endothelial cells. (A) Transcription factor activation screen (TF Activation Profiling Plate Array I - Signosis) of hUVEC 1 d after treatment (30 min) with either vehicle or 3 μg/mL Epirubicin was quanfified using an Omega Plate reader (BMG Labtech). As per manufacturer’s instructions, only transcription factors outside ± 10% of the blank sample and with a fold change > 2 compared to control samples are considered significant, and as such only TF obeying that criteria are displayed; (B) Transcript levels for key endothelial cell function factors, VEGF and eNOS, were quantified at baseline and after 1 d and 2 d after exposure to Epirubicin; data represents average fold change ± SE; Statistical significance was assessed by Two-Way ANOVA with Bonferroni Correction, and *p<0.05, **p<0.01, ***p<0.001, n=2.

